# ‘4-Aminoquinazoline-6, 7-diol’ derivatives for enhanced EGFR binding (as inhibitor) against Lung cancer

**DOI:** 10.1101/2022.02.14.480387

**Authors:** Pratibha Preeti Maurya

## Abstract

The Epidermal growth factor receptor (EGFR) have been considered as potential targets for cancer therapy due to their role in regulating proliferation and survival of cancer cells. In the present study, the four ‘4-aminoquinazoline-6, 7-diol’ derivative molecules were selected to the theoretically investigate their inhibitory activity against EGFR protein. Molecules were evaluated on the basis of: (i) in-silico cytotoxic assay on human cancer cell lines expressing EGFR (A549 and A431), (ii) in-silico kinase assay, (iii) impact of molecular interaction on EGFR, (iv) molecular interaction stability, (v) inhibitory impact of molecules on combined expression of EGF-EGFR and (v) ADME observations. Studies were observed on the scale of two known EGFR inhibitors. In-silico cytotoxicity screening results demonstrated that the A431 lung cancer cell line was vulnerable to compound abc1-4. However, A549 cells were less sensitive to our designed molecules. From kinase inhibition assay results, compound abc1&2 were found to be an inhibitor against EGFR. According to the molecular interaction analysis, compounds abc1-4 performed in similar fashion of positive controls, acting at the catalytic site of EGFR. Altogether, these potent compounds, with compromised bioavailability, could likely be developed as a promising EGFR targeted drug(s) for cancer therapy.

## INTRODUCTION

Cancer, a group of diseases characterized by uncontrolled growth and spread of abnormal cells, is the second leading cause of mortality worldwide. Among several types of cancer, lung cancer is at the leading cause of cancer-related death globally. Epidermal growth factor receptor (EGFR) is the well-known drug target for treating lung cancer, since it plays a major role in regulating proliferation and survival of cancer cells.

Various cell line are used for the study of EGFR over expressed cancers. In case of lung cancer, A549 & A431 are well known for these studies (*Suman et al., 2020*). EGFR, a member of the ErbB family of receptor tyrosine kinases, is composed of an extracellular domain, a single hydrophobic transmembrane region, an intracellular tyrosine kinase (TK) domain, and a C-terminal tyrosine-rich region (*Yang et al., 1998*). The EGFR signalling pathways are triggered by the specific growth factors (such as EGF and TGFα) to the extracellular domain, activating autophosphorylation and subsequently initiating downstream signalling pathways, responsible for cell proliferation, survival, differentiation and apoptosis evasion. Importantly, more than 60% of non-small cell lung cancer have been involved in the overexpression of EGFR; therefore, targeting this protein is an important strategy for lung cancer treatment. To date, several EGFR inhibitors have been reported such as Erlotinib and Gefitinib (*Singh et al., 2016*). Among them, Erlotinib has been the first-line drug for lung cancer patients harboring wild-type EGFR (IC50 against EGFR is 2.6 nM). However, these drugs have several side effects, e.g., anemia, balance impairment and dizziness. Moreover, acquired drug resistances of and Erlotinib (T790M) have been detected after 1.5 year treatment. Accordingly, discovery of novel EGFR inhibitors is critically needed.

EGFR is a tyrosine kinase receptor that is widely overexpressed in cancers such non-small-cell lung cancer, metastatic colorectal cancer, glioblastoma, head and neck cancer, pancreatic cancer, and breast cancer. Common mutations and truncations to its extracellular domain, such as the EGFR viii truncations, as well as to its kinase domain, such as the L858R and T790M mutations, or the exon 19 truncation, all contribute to increased EGFR activity. RAS-RAF-MEK-ERK MAPK and AKT-PI3K-mTOR are two downstream pro-oncogenic signalling pathways activated by these EGFR mutations. These pathways then activate a variety of biological outputs that aid cancer cell proliferation, including persistent initiation and advancement through the cell cycle. The molecular mechanisms that regulate EGFR signal transduction, including EGFR structure and mutations, ligand binding and EGFR dimerization, and the signalling pathways that lead to G1 cell cycle advancement, are discussed here. EGFR signalling pathways are studied for stimulation of CYCLIN D expression, activation of CDK4/6, and inhibition of cyclin-dependent kinase inhibitor proteins (CDKi). We also go over the benefits and drawbacks of EGFR-targeted therapy, as well as the potential for combining them with CDK4/6 inhibitors (*Prabhu et al., 2018*).

In recent years, 4-anilinoquinazolines have emerged as a versatile template for inhibition of a diverse range of receptor tyrosine kinases. The most widely studied of these is the epidermal growth factor receptor (EGFR), with the small-molecule inhibitor Gefitinib being the first agent from this class to be approved for the treatment of non-small cell lung cancer refractory to prior chemotherapeutic intervention. Subsequent research aimed at further exploration of the SAR of this novel template has led to discovery of highly selective compounds that target EGFR (*Barbosa et al., 2014*). These compounds act via competing with ATP for binding at the catalytic domain of tyrosine kinase. Later on, a great structural variety of compounds of structurally diverse classes have proved to be highly potent and selective ATP-competitive inhibitors. Among them, 4-anilinoquinoline-3-carbonitriles and others provide the necessary binding properties for inhibition of the ErbB family of tyrosine as they mimic the adenine portion of ATP. In this study, we present a new sub-family of compounds containing 4-aminoquinazoline, and Quinoline core as promising potent and selective EGFR inhibitors. Our strategy is directed toward designing a variety of ligands with diverse chemical properties hypothesizing that the potency of these molecules might be enhanced by adding alternative binding group such as sulphonamide and carboxylic group in the aniline moiety. In this way, such substitution pattern could target different regions of the ATP-binding site of the protein kinase domain to create differentially selective molecules. The design of our ligands was done based on previous structure–activity relationship (SAR) of 4-anilinoquinazolines and earlier work with Quinazoline-based inhibitors of EGFR which established that 6, 7-dialkoxy substitution is compatible with good activity, and pivotal interactions between the receptor and the inhibitors (*Barbosa et al., 2014*). In more recent approach, it was found that dual inhibition of EGFR and ErbB-2 may offer increased activity over agents which target only one of these receptor kinases. After discovery of Lapatinib, it was claimed that 4 position of the aniline can tolerate a lot of bulky substituents; this leads to fundamental change in the pharmacophore and claims moderate ErbB-2 activity with little or no EGFR activity. In this direction and in an approach to enhance the selectivity toward ErbB-2, we introduced larger moiety at 4 position of the aniline such as heterocyclylsulfonamido moiety in a fashion similar to Lapatinib which binds in the ATP-binding cleft, so that the bulky group could be oriented deep in the back of the ATP binding site and makes predominantly hydrophobic interactions with the protein mimicking the aniline derivative group of Lapatinib (*Wilson et al., 2015*). Moreover, in two compounds we replaced the aniline moiety with substituted piperazine fragment to assess if such dramatic change in the pharmacophoric model of anilinoquinolines or quinazoline will have appreciable effect on the activity. The binding mode and docking energy of the designed compounds in EGFR homology model could be helpful tool for predicting their mechanism of antitumor activity of the target compounds.

## MATERIALS AND METHODS

The present study is intended to computationally designing as well as evaluation of EGFR inhibitors. Molecules were designed on the basis of evolutionary repositioning of fragments from known EGFR binders. Aim was to re-define the molecules with more molecular interactions. At the very first step, cytotoxic activity was evaluated through two lung cancer cell line models for A549 & A431 (*Suman et al., 2020*). Two known EGFR binder were considered as positive controls. Consistency of molecular interaction, of designed molecules & known EGFR binders, was evaluated with in-silico kinase assay model (*Wagner et al., 2021*). Molecular interaction at atomic level was observed through molecular docking studies, with PDB 4i23 (*Gajiwala et al., 2013*) and AutoDock Vina (*Trott & Olson, 2010*). Furthermore, molecular interaction stability at atomic level was deeply observed through GROMACS based Molecular Dynamics studies. Impact of molecules was observed in extrapolation through existing system model for normal EGFR. It was used for evaluation of Inhibitory impact of molecule on interaction of EGF-EGFR, under the influence of other components of system (*Bidkhori et al., 2012*). Designed molecules were finally processed for evaluation of bioavailability through SwissADME (*Daina et al., 2017*).

## RESULTS

### Designing of molecules

Compound ‘4-aminoquinazoline-6, 7-diol’ derivatives (abc1-4) were designed on the basis of repositioning of fragments from existing potential molecules (Figure 2). Four molecules showed the possible enhanced potency over existing ones. Here designed molecules were evaluated for their molecular performance. Two positive controls i.e. known EGFR binder, were used: first one was Erlotinib & second one was co-crystallised ligand at PDB: 4i23. Molecules were evaluated on the basis of: (i) in-silico cytotoxic assay, (ii) in-silico kinase assay, (iii) impact of molecule on EGFR, (iv) molecular interaction stability, (v) inhibitory impact of molecule on interaction of EGF-EGFR and (v) ADME observations.

**Figure 1.**
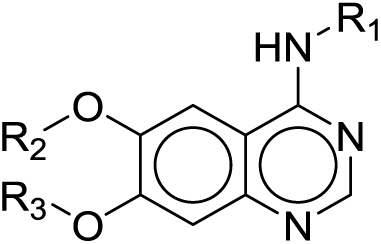
Based scaffold (4-aminoquinazoline-6,7-diol) for EGFR binding.

**Figure 2.**
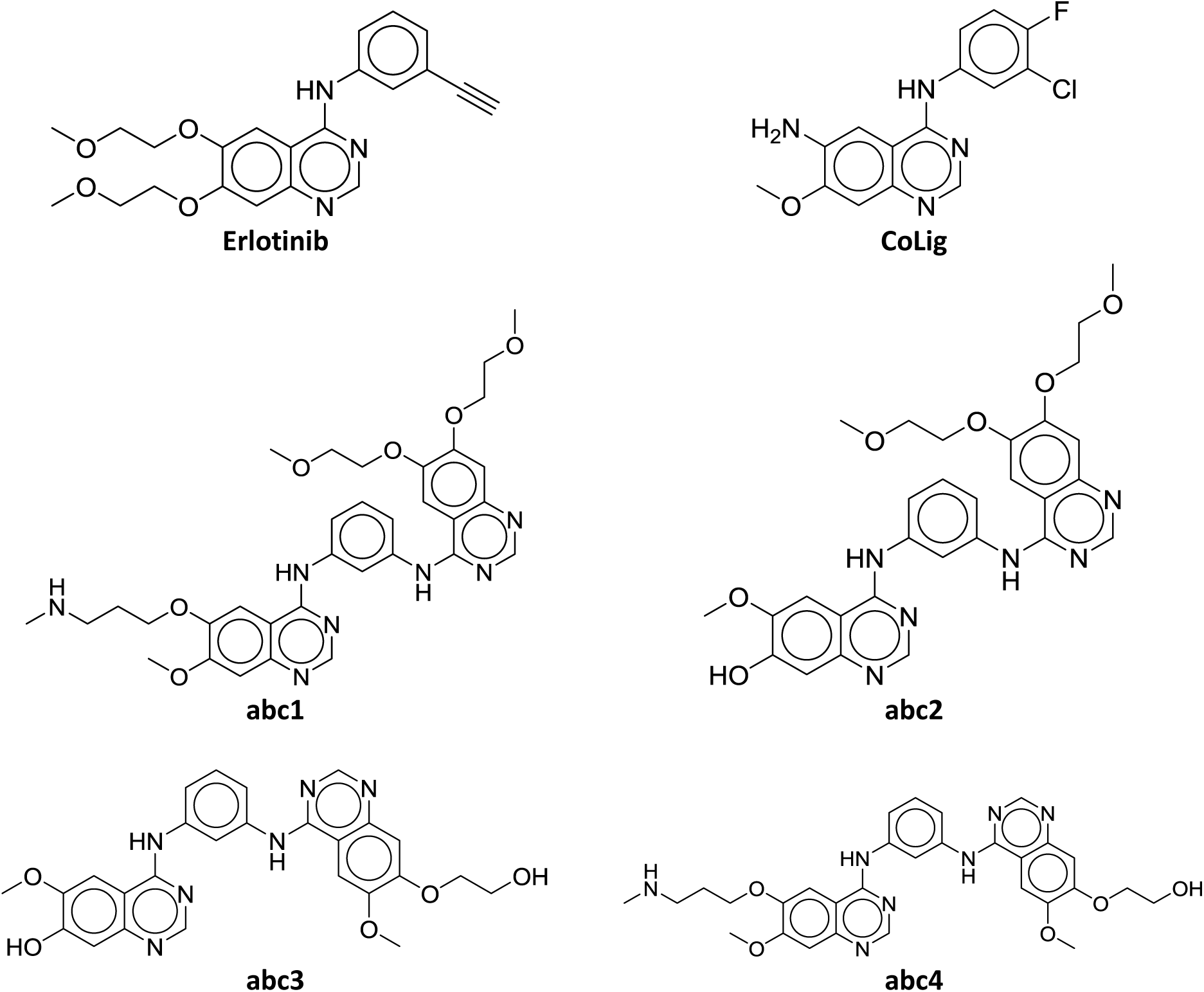
Known EGFR inhibitors & Designed molecules.

### Cytotoxicity

Initially, abc1-4 and known drugs (Erlotinib & CoLig) were subjected to in-silico cytotoxicity screening against A549 and A431 cell lines expressing EGFR. The results revealed that the A431 lung cancer cell line was vulnerable to compounds abc1-4. However, A549 cells was less sensitive to the derivatives. These four compounds were evaluated through their half maximal inhibitory concentration (IC50) values. The IC50 values of abc1-4 and the known drugs towards the two cancer cell lines are summarized in Table 1. The results revealed that in the case of EGFR-expressing cells, the anti-lung cancer potential of derivatives ranges between IC50 of 7.1 µM to 8.0 µM for A549; and between 2.2 µM to 3.4 µM for A431. In addition, It was found that the IC50 value of the A431 cell line treated with compound abc1&4 (IC50 of approximately 2.5 µM) was similar to CoLig (IC50 of approximately 2.2 µM), while for compound abc2&3 (IC50 of approximately 3.3 µM) was similar to Erlotinib (IC50 of approximately 3.0 µM). Similarly in case of A549 cell line, it was found that performance of compound abc1-4 (IC50 of approximately 7.0 µM) was similar to Erlotininb (IC50 of approximately 7.9 µM). Furthermore, these compounds were processed for in-silico kinase inhibitory activity assay against the EGFR-Kinase.

**Table 1.** Cytotoxic performance of molecules

| Molecules | IC<sub>50</sub> nM against A549 (Active-Inactive class values ranging between 1-0) | IC<sub>50</sub> nM against A431 (Active-Inactive class values ranging between 1-0) |
| --- | --- | --- |
| Erlotinib | 7926.6 (0.677) | 3007.65 (1.099) |
| CoLig | 9742.4 (0.275617) | 2244.2 (0.5426) |
| abc1 | 7178.0 (0.4955) | 2516.88 (0.711826) |
| abc2 | 7184.95 (0.501696) | 3304.99 (1.084) |
| abc3 | 7160.75 (0.558) | 3359.76 (1.01457) |
| abc4 | 7160.4 (0.5250) | 2553.72 (0.681155) |

### Kinase inhibition

To further evaluate that compounds abc1-4 can inhibit the EGFR-TK proteins, the kinase inhibitory activity assays were conducted in comparison to the known inhibitors (erlotinib). As shown in Table 2, the kinase inhibitory score against A431 of compounds were: Erlotinib (1.5006), CoLig (1.441882), abc1 (1.700237), abc2 (2.310034), abc3 (0.078865), abc4 (−0.24669). Results suggested the potency of abc1 & abc2 against EGFR. Here, kinase inhibitory activity in cell line A431 was represented by ‘C0_K1/C1_K0’ with higher values as positive controls.

**Table 2.**
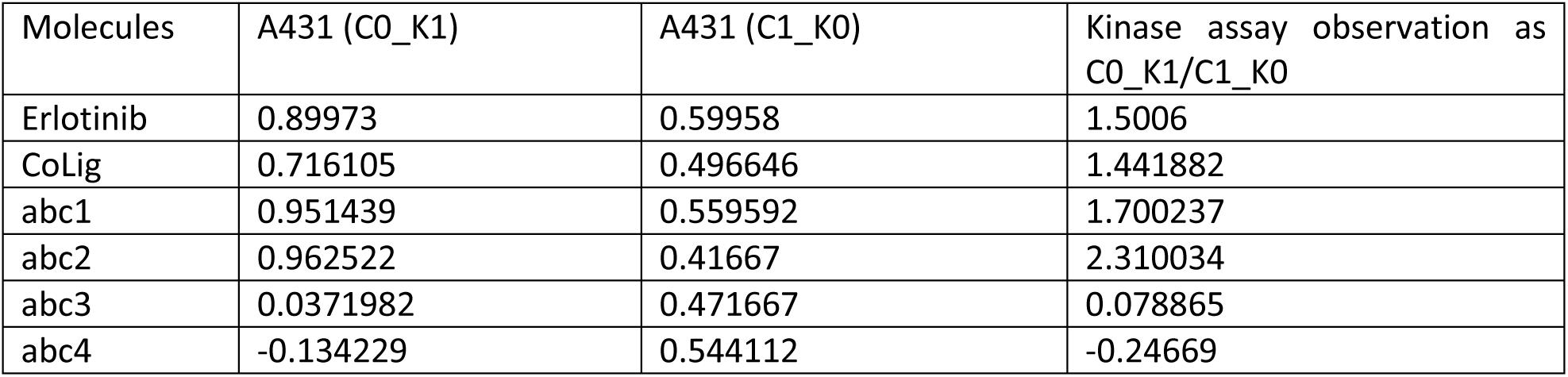
Kinase assay output for tested compounds

### Impact of molecules on EGFR using molecular docking

Molecular docking was conducted to investigate the binding mechanism of the focused derivatives towards EGFR-TK in comparison to the known drugs Erlotinib & CoLig. It was found that the derivative EGFR-TK complexes gave a binding affinity approximately similar or higher than the Erlotinib & CoLig-EGFR complex. The binding affinity and residual patterns of underlying interactions of all focused ligands are shown in Table 3 and Figure 3. In the case of Erlotinib complexed with EGFR-TK, its residue-718 & 793 were hydrogen bounded. While main H-bond interaction was found with residue-745. All derivatives were consistently bear H-bond with residue-745. ‘abc2’ & ‘abc3’ showed additional interaction with residue-793 as Erlotinib. Besides these, residue 720 & 724 were also found to be involved in interaction with ligands.

**Table 3.**
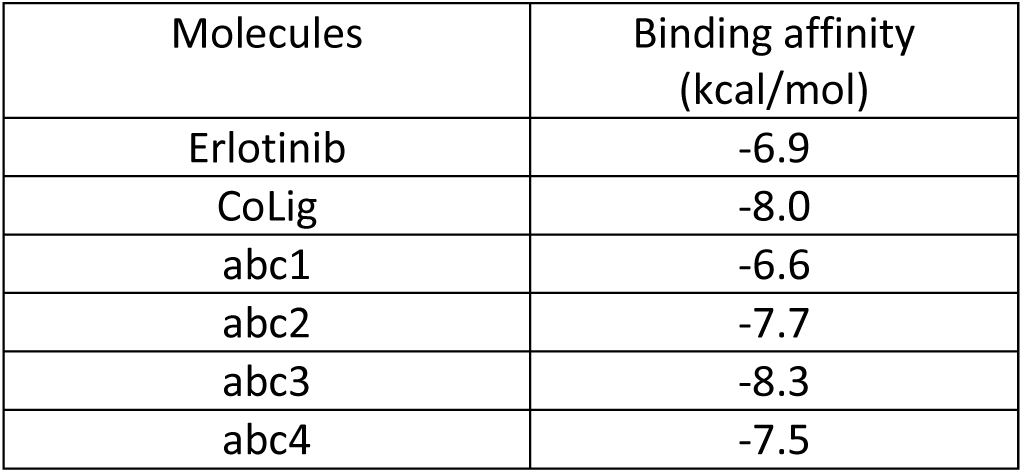
Binding affinity of ligand

| Molecules | Binding affinity (kcal/mol) |
| --- | --- |
| Erlotinib | -6.9 |
| CoLig | -8.0 |
| abc1 | -6.6 |
| abc2 | -7.7 |
| abc3 | -8.3 |
| abc4 | -7.5 |

**Figure 3.**
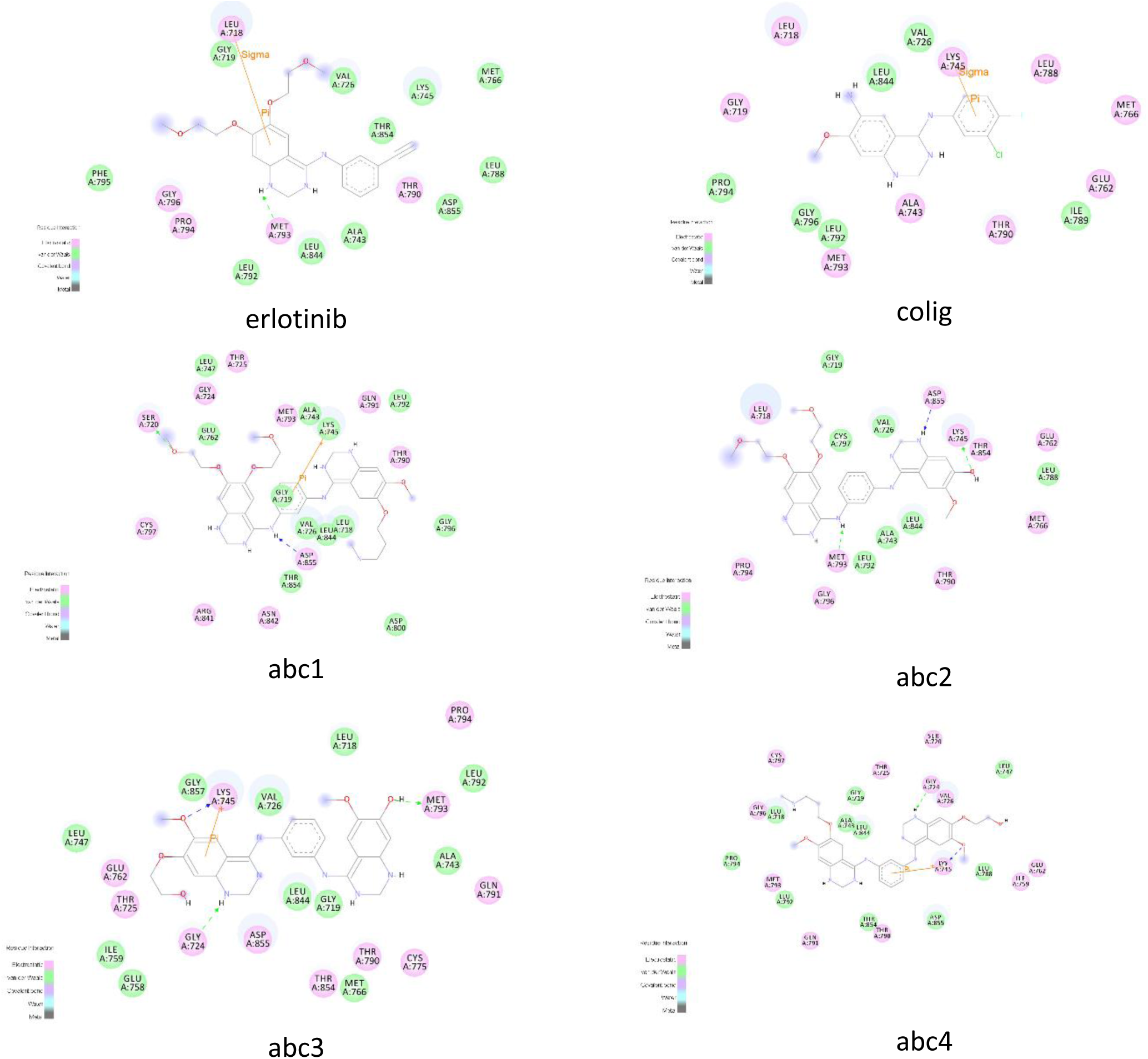
Molecular interactions between ligands & PDB 4i23.

These results were further used to investigate the pattern of impact of ligands on EGFR. Docking force on protein by ligand due to protein-ligand interaction was observed as: lower limit was defined by the positive control compounds as 0.2436. Compounds with value greater than 0.2436, were considered as more significant which showed required impact on the EGFR for inhibition. In reference of Erlotininb, It was found that all four abc1-4 showed required force on EGFR for inhibition. While in reference of CoLig, abc2 & 4 showed potential impact (Table 4, Figure 4).

**Table 4.**
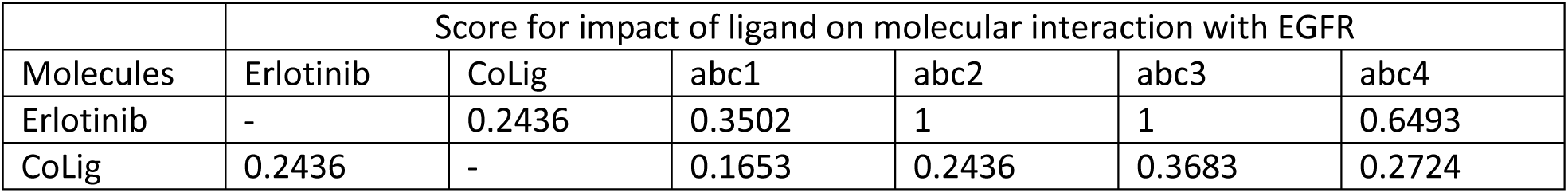
Impact of ligand on molecular interaction with EGFR

**Figure 4.**
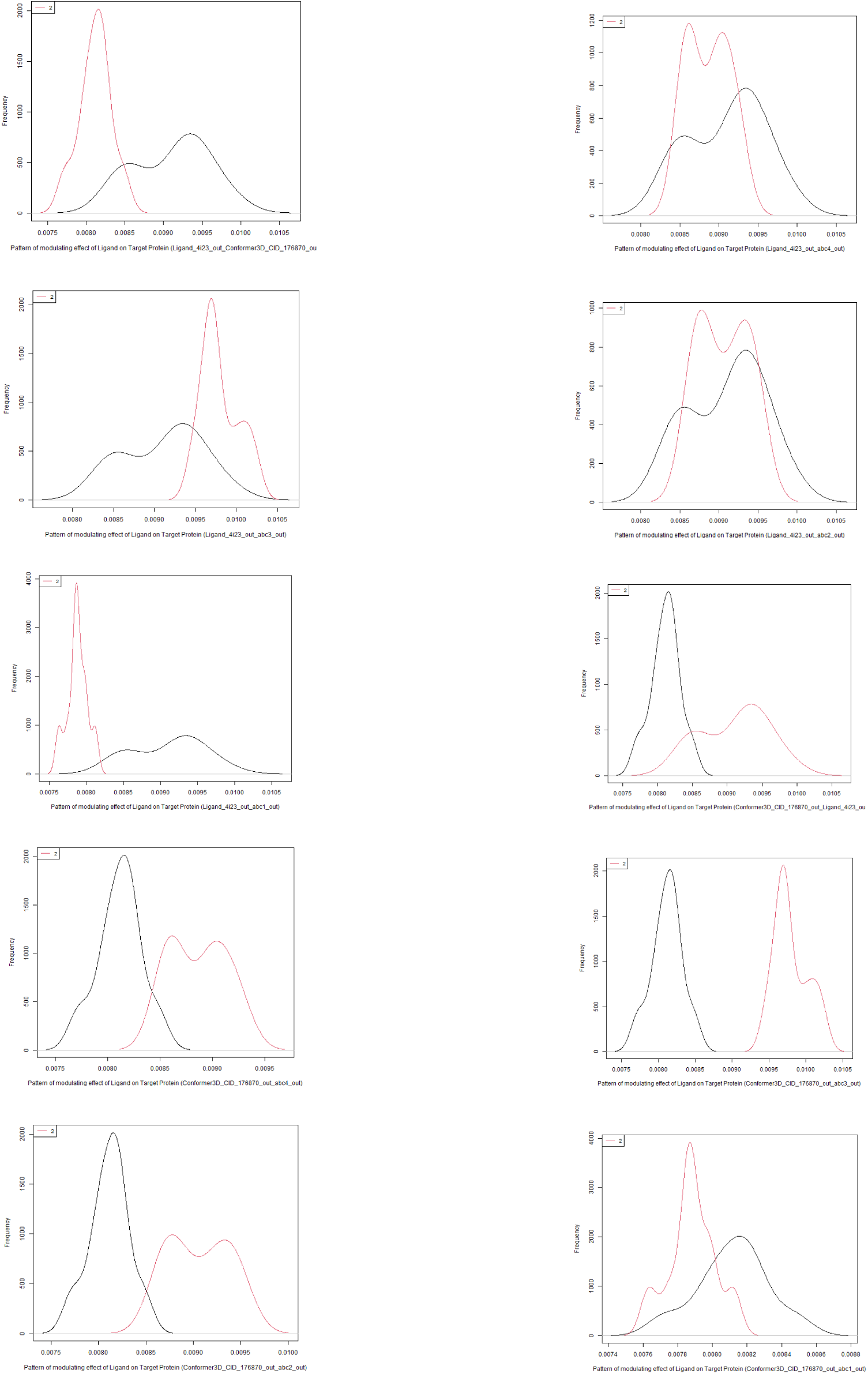
Impact of ligands on EGFR.

### Initial interaction impact of molecules on the stability of EGFR

Initial interaction impact of molecules on the stability of EGFR was observed through molecular dynamics of first 1000 ps. Initial fluctuation was observed in terms of average RMSF value during 1000 ps (Figure 5 & 6). Initial fluctuations define the relative behaviour of molecules towards the target protein. It was observed that none of the designed molecule was as stable as the Erlotinib & CoLig; but abc1 & abc2 behaved in similar fashion. While abc3 was found to be most stable. Molecule abc4 was found to be most fluctuating.

**Figure 5.**
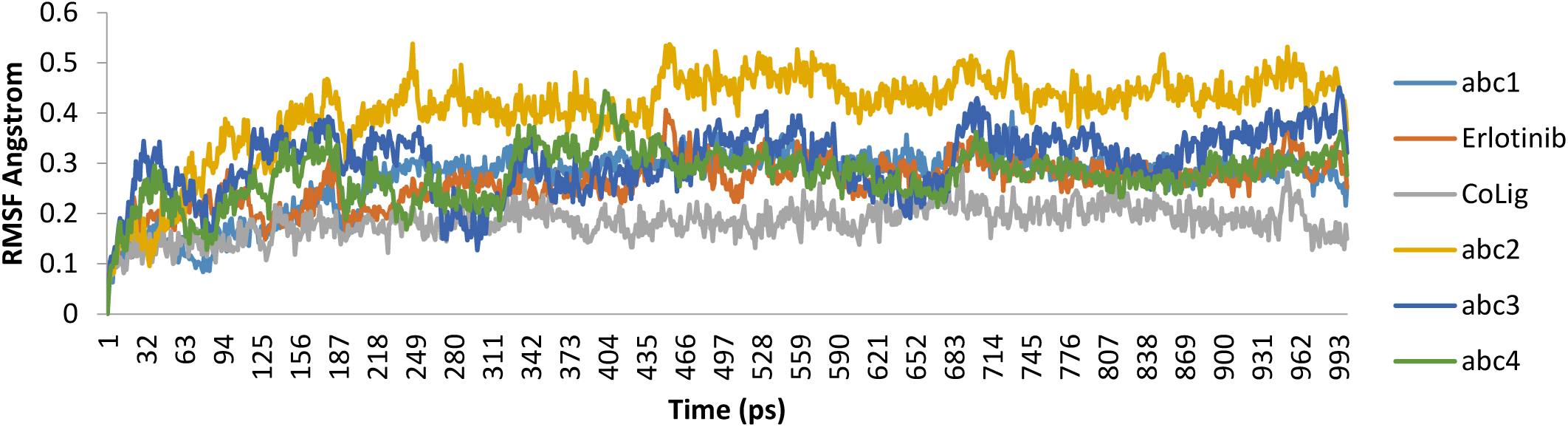
RMSF found on MD simulation for 1000 ps.

**Figure 6.**
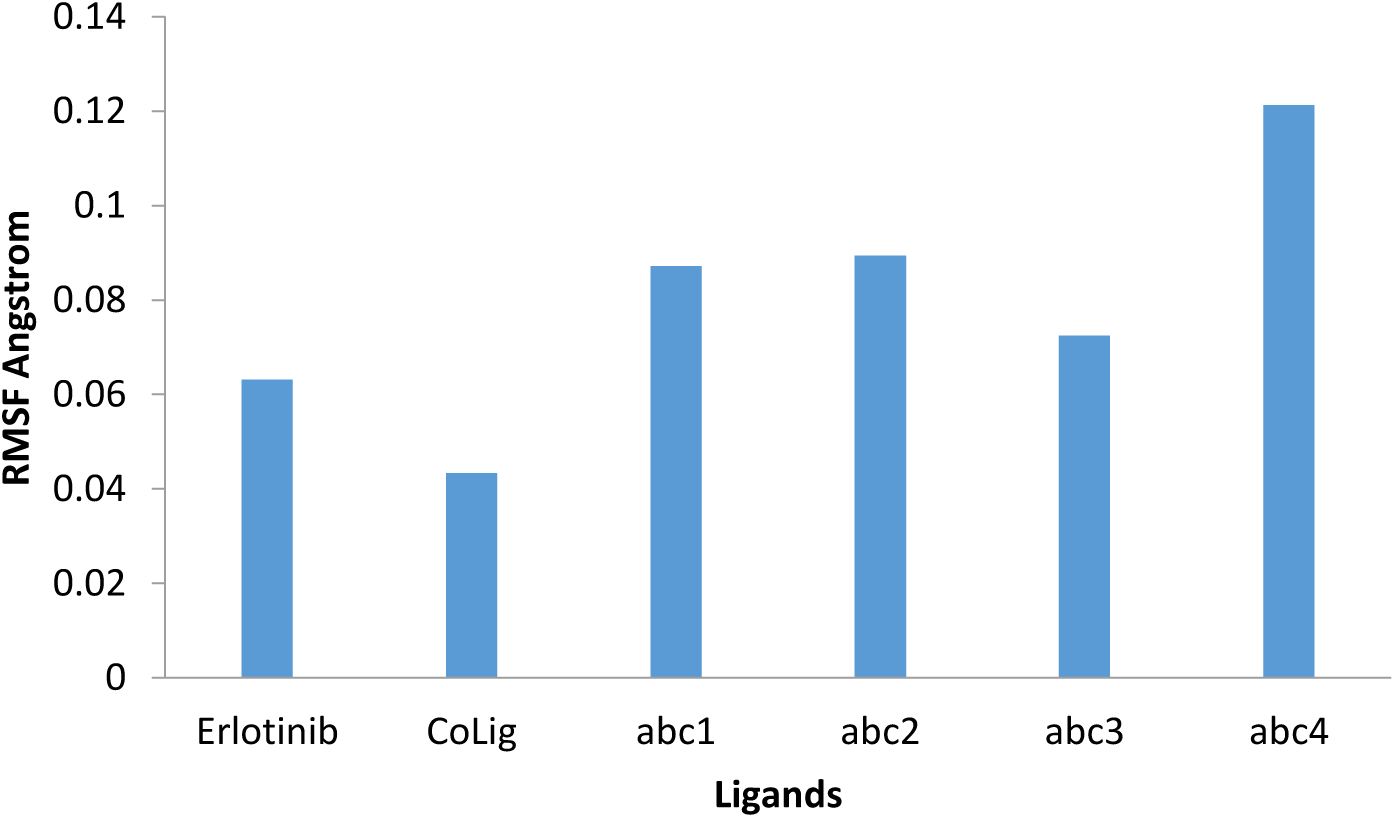
Relative RMSF of ligands.

### Inhibitory impact of molecule on interaction of EGF-EGFR, under the influence of other components of system

This was performed through system model. Normal EGFR-system model was borrowed, and cancer EGFR-system model was developed. Treatment of compounds abc1-4 & controls were observed with concentration of 1-10 µM. Default k3 value, of normal EGFR system, was 2.014. It was decreased in sequencially manner to increase the expression of EGF-EGFR. At the point of k3 value with highest expression of EGF-EGFR was considered as EGFR cancer system model. Firstly ligand concentration of 10 µM was used for evaluating the impact of treatment. Derivative abc1 & abc4 were found to be most potential for inhibition og EGF-EGFR expression. Furthermore, Ligand concentration gradient of 1 to 10 µM was applied with normal as well as cancer EGFR model. Gradient tratement also showed same bahivour of derives abc1 & abc4 (Table 5, Figure 7 & 8).

**Table 5.** Treatment of 10 µM ligands at system model

| Molecules | Binding affinity (kcal/mol) | ki (nM) | Model Cell line without treatment | Cell line model coefficient (Normal) | Cell line model coefficient (cancer) | Model Cell line treated with molecules at 10 μM |
| --- | --- | --- | --- | --- | --- | --- |
| Erlotinib | 6.9 | 114227.7 | 1.087544449 | 2.014 | 0.5 | 0.543772224 |
| CoLig | 8 | 731278.4 | 1.013674682 | 2.014 | 0.5 | 0.506837341 |
| abc1 | 6.6 | 68844.35 | 1.145255209 | 2.014 | 0.5 | 0.572627604 |
| abc2 | 7.7 | 440737.2 | 1.02268926 | 2.014 | 0.5 | 0.51134463 |
| abc3 | 8.3 | 1213349 | 1.008241649 | 2.014 | 0.5 | 0.504120825 |
| abc4 | 7.5 | 314468.8 | 1.031799655 | 2.014 | 0.5 | 0.515899827 |

**Figure 7.**
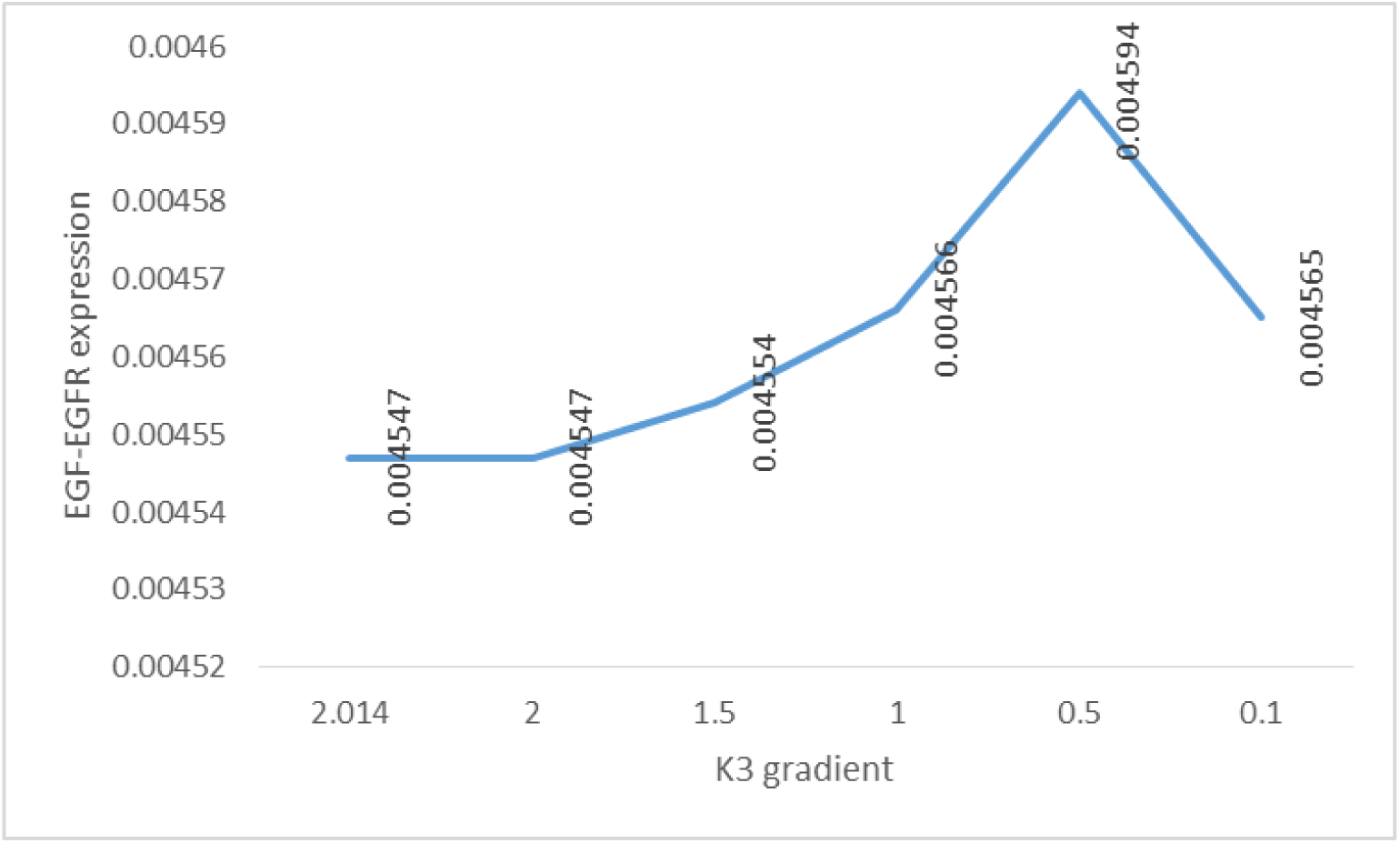
Preparation of cancer EGFR-system model.

Performance of ligand was observed with concentration gradient of 01-10 micro Molar. Molecule ‘abc1’ performed better than Erlotinib. Molecules abc2 & 4 performed in approximately similar fashion, but not better than Erlotinib (Table 6, Figure 9).

**Table 6.** Treatment with concentration gradient of 1-10 µM

|  | Model Cell line treated with molecules of concentration of |  |  |  |  |  |
| --- | --- | --- | --- | --- | --- | --- |
| Molecules | 1 μM | 2 μM | 4 μM | 6 μM | 8 μM | 10 μM |
| Erlo | 0.004594 | 0.004593 | 0.004593 | 0.004592 | 0.004591 | 0.004591 |
| CoLig | 0.004594 | 0.004594 | 0.004594 | 0.004594 | 0.004594 | 0.004594 |
| abc1 | 0.004593 | 0.004593 | 0.004592 | 0.004592 | 0.004589 | 0.004588 |
| abc2 | 0.004594 | 0.004594 | 0.004594 | 0.004594 | 0.004593 | 0.004593 |
| abc3 | 0.004594 | 0.004594 | 0.004594 | 0.004594 | 0.004594 | 0.004594 |
| abc4 | 0.004594 | 0.004594 | 0.004594 | 0.004593 | 0.004593 | 0.004593 |

**Figure 8.**
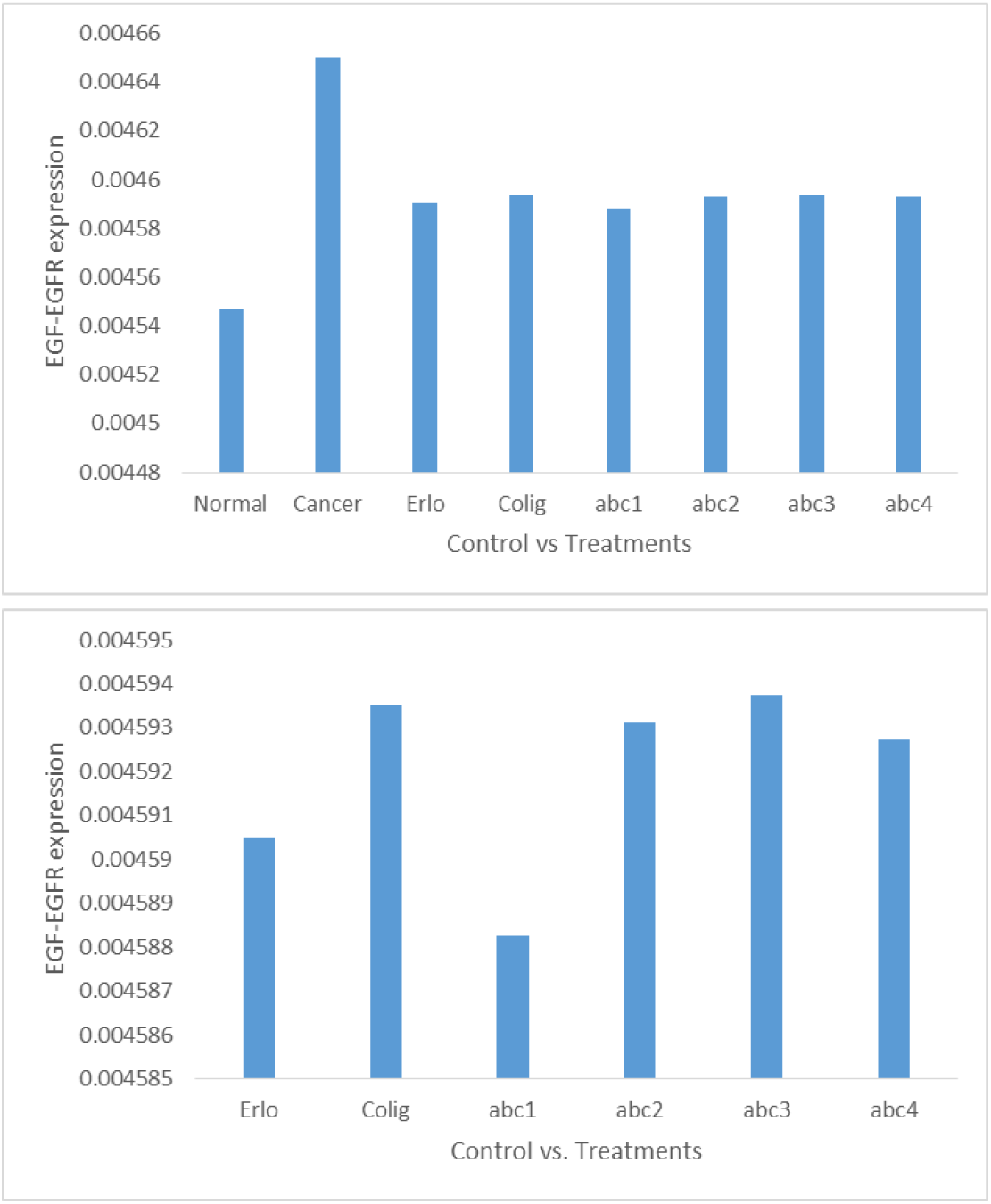
EGF-EGFR expression in Normal-Cancer model with Control & Treatments.

**Figure 9.**
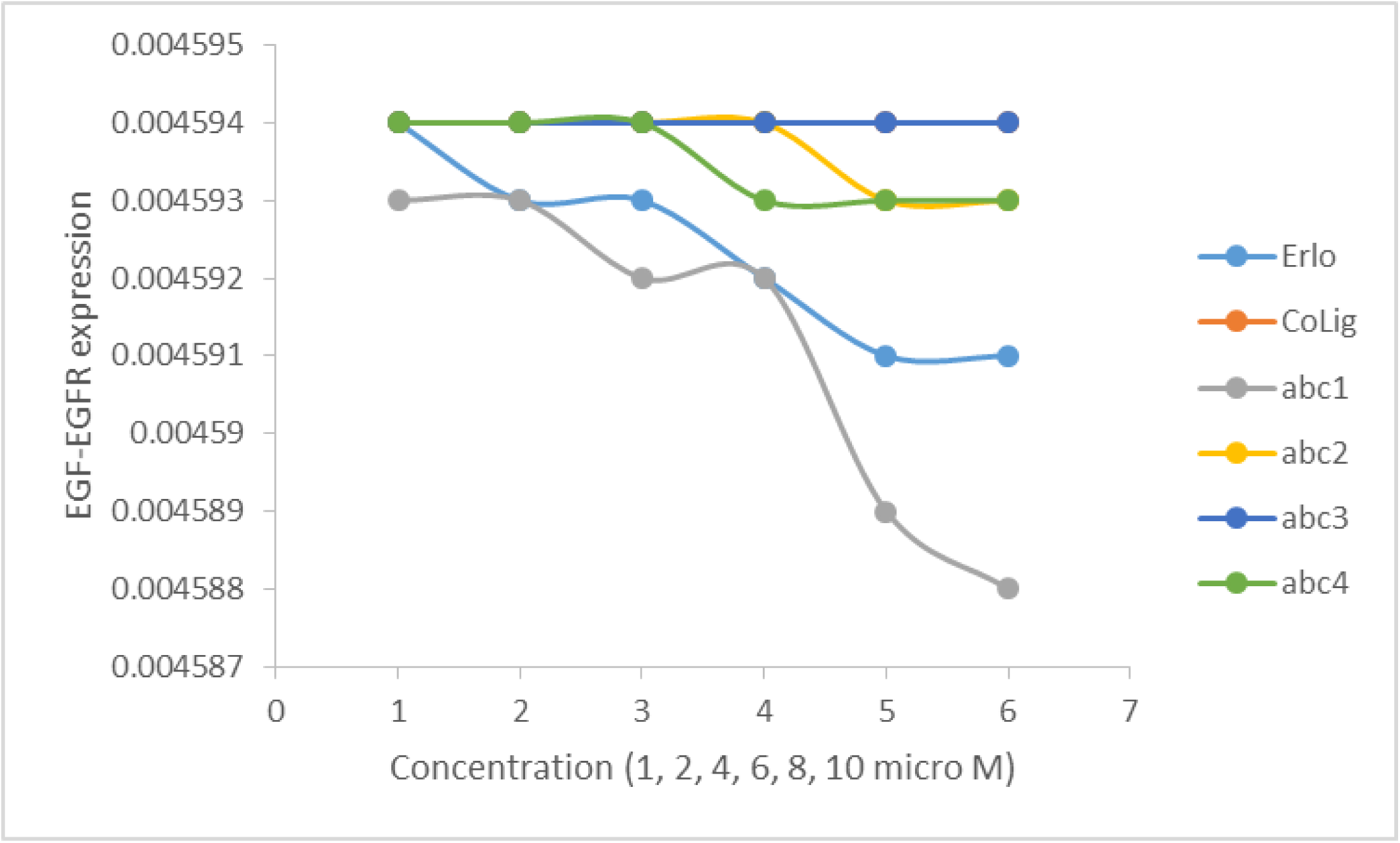
Performance of Ligands.

### ADME observation

Overall aromatic heavy atoms, rotatable bonds and Hydrogen bond interactions were increased in new compounds. But due to increase in TPSA, new compounds are not accessible to BBB. Solubility also decreased in comparison of reference compounds. Synthetic accessibility increased. In this way, overall two newly designed molecules showed enhance target activity, but with compromised bio-availability than reference control compounds.

## DISCUSSION

The 4-aminoquinazoline have been previously reported to exhibit anti-tumor activities through binding EGFR against several types of cancer. Therefore, we expected that newly designed 4-aminoquinazoline derivative can inhibit EGFR-TK activity. Considering the data from the in silico cytotoxic activity of the four 4-aminoquinazoline derivatives against lung cancer cell lines, we found that compounds were more susceptible to EGFR-expressing cell lines, A431 lung cancer cell line, whereas A549 cells were comparatively less sensitive to these derivatives. In addition, designed compounds showed cytotoxicity to A431 higher than A549. This is because (i) the EGFR expression level found in A431 cells is dramatically higher than that found in A549 and (ii) A549 cells exhibits KRAS mutation, which constitutively activates downstream MAPK signalling pathways, causing a compensatory mechanism. The cytotoxic effect on cancer cell lines of these 4-aminoquinazoline derivatives was due to their inhibitory activity against EGFR-TK, in a manner similar to the known inhibitors. In addition, both these derivatives showed approximately similar IC50 than the known drugs, suggesting that these 4-aminoquinazoline derivatives could be safe similar to the known drugs. However, the results from the cell-based assay were inconsistent with kinase inhibition in which compounds abc1 and abc2 were potent against kinase, whereas compounds abc3 & abc4 showed less inhibitory activity towards kinase. This is because cells may have several factors involved in cell growth inhibition such as cell permeability and compound degradation within the cell. Therefore, compounds with good inhibitory activity from both kinase inhibition and cell-based inhibition were selected marked for key attention to study the binding patterns at the atomic level.

Although the docking binding affinity compounds abc2 and abc4 in complex with EGFR was found approximately same than that of the known inhibitors; their ligand–protein interactions were also similar. It was found that all the ligands including controls showed consistency in interaction with two residues LYS745 & MET793. Besides these, derived showed more interaction with other residues. Overall, derivates abc1 & abc4 were observed to be more consistent with known EGFR inhibitors. Molecular interaction stability analysis showed that, all the derivative are less stable than known drug at the level of molecular interaction. Among the derivatives, abc3 is most stable, abc4 is least stable, while abc1 & abc2 were found to approximately similar in stability.

Inhibitory impact of molecule on interaction of EGF-EGFR, under the influence of other components of system was accomplished using the system model. The normal EGFR-system model was adapted, and a cancer EGFR-system model was created. Compounds abc1-4 and controls were treated with concentrations ranging from 1 to 10 µM. The normal EGFR system’s default k3 value was 2.014. It was gradually reduced in order to increase the expression of EGF-EGFR. The EGFR cancer system model was chosen at the k3 value with the highest expression of EGF-EGFR. To begin, a ligand concentration of 10 µM was used to assess the impact of treatment. The derivatives abc1 and abc4 were discovered to have the greatest potential for inhibiting EGF-EGFR expression. Furthermore, a ligand concentration gradient of 1 to 10 µM was used with both the normal and cancer EGFR models. Gradient treatment also demonstrated the same behaviour of derives abc1 and abc4.

Finally ADME study showed that in novel compounds, the number of aromatic heavy atoms, rotatable bonds, and hydrogen bond interactions all increased. However, novel compounds are not accessible to BBB because to an increase in TPSA. In compared to reference compounds, solubility also reduced. The availability of synthetics has improved. In this fashion, two newly developed molecules outperformed reference control compounds in terms of target activity, but at the expense of bioavailability.

## CONCLUSIONS

In this study, we implemented computational studies to identify novel EGFR inhibitors. In-silico cytotoxicity screening results showed that the A431 lung cancer cell line was vulnerable to compound abc1-4. The cytotoxic effect on cancer cell lines of these derivatives was due to their inhibitory activity against EGFR. From binding interaction analysis, it was found that compounds abc1-4 are pounding sufficient force on protein EGFR. Our study successfully identified advanced EGFR inhibitors abc1 & abc4, which can be used as promising starting points for subsequent drug discovery programs against lung cancer.

## Acknowledgement

Author is thankful to Computational Biology for Biochemical Experiments (CBBE), Lucknow, India for providing research facilities.

